# Win-paired cues modulate the effect of dopamine neuron sensitization on decision making and cocaine self-administration: divergent effects across sex

**DOI:** 10.1101/2023.05.19.541413

**Authors:** Tristan J. Hynes, Chloe S. Chernoff, Kelly M. Hrelja, Maric T.L. Tse, Dimitrios Avramidis, Melanie R. Lysenko-Martin, Lucas Calderhead, Sukhbir Kaur, Stan B. Floresco, Catharine A. Winstanley

**Author notes:** Correspondence should be addressed to Dr. Tristan J. Hynes or Dr. Catharine A. Winstanley Address: Djavad Mowafaghian Centre for Brain Health, Department of Psychology, 2215, Wesbrook Mall Vancouver, BC, V6T 1Z3, Canada. Department of Psychology, University of Cambridge, Cambridge, United Kingdom (Tristan J. Hynes, Catharine A. Winstanley). These two authors made equal contributions (Tristan J. Hynes, Chloe S. Chernoff).

## Abstract

Psychostimulant use and engagement with probabilistic schedules of reward both sensitize the mesocorticolimbic dopamine system. Such behaviours may act synergistically to explain the high comorbidity between stimulant use and gambling disorder. The salient audiovisual stimuli of modern electronic gambling may exacerbate the situation. To probe these interactions, we sensitized ventral tegmental area (VTA) dopamine neurons via chronic chemogenetic stimulation while rats learned the rat gambling task in the presence or absence of casino-like cues. The same rats then learned to self-administer cocaine. In a separate cohort, we confirmed that our chemogenetic methods sensitized the locomotor response to cocaine, and potentiated phasic excitability of VTA dopamine neurons through in vivo electrophysiological recordings. In the absence of cues, sensitization promoted risk-taking in both sexes. When rewards were cued, sensitization expedited the development of a risk-preferring phenotype in males, while attenuating cue-induced risk-taking in females. While these results provide further confirmation that VTA dopamine neurons critically modulate risky decision making, they also reveal stark sex differences in the decisional impact which dopaminergic signals exert when winning outcomes are cued. As previously observed, risky decision-making on the cued rGT increased as both males and females learned to self-administer cocaine. The combination of dopamine sensitization and win-paired cues while gambling lead to significantly greater cocaine-taking, but these rats did not show any increase in risky choice as a result. Cocaine and heavily-cued gambles may therefore partially substitute for each other once the dopamine system is rendered labile through sensitization, compounding addiction risk across modalities.

## Introduction

Gambling and substance use disorders are highly comorbid, with cocaine being the most commonly abused substance among people with gambling disorder [1–4], potentially indicative of common brain substrates across these addictions [5, 6]. Substantial human and preclinical evidence point to the dopamine system as being heavily involved in the transition to addiction, whether it be to gambling [7] or to drugs [8, 9]. The dopamine system is also responsible for mediating the biobehavioural response to reward cues, and indeed, cues play a vital role in both gambling and cocaine addiction [10–12]. Much like the cues associated with cocaine (e.g., a syringe or crackpipe) evoke powerful urges to take the drug, the exciting lights and sounds of the modern casino environment can drive the urge to gamble [13].

Repeated administration of cocaine results in psychomotor sensitization through neuroplastic processes within the mesolimbic dopamine system [14, 15]. Behaviourally, this is typically measured in locomotor assays, in which rats become increasingly hyperactive when cocaine is administered. However, sensitization also occurs to the ability of reward-paired cues to act as conditioned reinforcers and motivate operant responding [16], a phenomenon known as incentive sensitization [17]. The use of cocaine may therefore potentiate the ability of audiovisual cues to influence engagement with electronic gambling machines.

Data from both humans and rats show that such cues increase risky decision making in simulated laboratory-based gambling tasks [18–20]. Greater risky decision making is associated with gambling disorder as well as cocaine use disorder [21, 22], and plays an important role in the addicted state [23–26]. Furthermore, rodent data suggest the act of engaging with probabilistic reinforcement schedules or cues that predict uncertain rewards can itself sensitise the locomotor and reinforcing effect of psychostimulants [27–30]. As such, the ability of cocaine to amplify the motivational influence of cues could drive problematic gambling as well as further drug use in a synergistic manner.

In support of this, we have previously shown that cocaine self-administration increases risky decision-making on the cued rGT, and this can be attenuated by chemogenetic inhibition of dopaminergic neurons within the ventral tegmental area (VTA) in both male and female rats [31, 32]. Dampening neurotransmission in this pathway during acquisition of the cued rGT also results in more optimal decision-making in males only [32]. Such data fit with the theory that win-concurrent cues and probabilistic rewards within the cued rGT can drive risky choice by eliciting dopamine release, and cocaine self-administration further sensitises the dopaminergic system and therefore exacerbates risky decision making [33]. However, the dopaminergic regulation of choice strategies in females may differ.

We therefore developed the experiments presented here with the following hypotheses in mind. If cocaine-taking promotes risky choice by sensitising the dopamine system, then facilitating the development of sensitisation should potentiate risky decision making. When reward-paired cues are combined with risky choice, the effects of dopamine sensitisation may be compounded, at least in males. Sensitised animals should also self-administer more cocaine, and such cocaine self-administration should produce a synergistic increase in risky decision making, particularly on the cued rGT. To this end, we used chemogenetics to repeatedly activate VTA dopamine neurons to evoke a biobehaviourally sensitized phenotype (i.e., increased locomotor reactivity to cocaine and phasic activity of VTA DA neurons). We then applied this sensitizing regimen of chemogenetic stimulation as separate animals learned to perform the uncued and cued rGT. We evaluated the response of chemogenetic sensitization of VTA DA neurons on the development of risk preference profiles, cocaine self-administration, and the influence of cocaine self-administration on decision making.

## Methods

### Subjects

All rats used for these experiments were bred in-house from wildtype female Long-Evans (Charles River, St. Constant, QC) and transgenic male rats that expressed Cre recombinase in neurons producing tyrosine hydroxylase (TH: Cre, 3.1Deis, RRRC #00659; Columbia, MO). Progeny of both sexes testing hemizygous for co-expression of Cre and TH (transgene positive; TG+) were used as experimental animals. Control groups were comprised of equal numbers of TG+ and TG-animals (see supplemental methods for further information).

### Viral infusion surgery

At PND 35, experimental animals received bilateral infusions (0.5 µL/hemisphere) of AAV5-hSyn-DIO-hM3D(Gq)-mCherry (7×10^12^ vg/mL; Addgene) into the VTA as described in [32]. TG+ controls received identical bilateral infusions of AAV5-hSyn-DIO-mCherry (7×10^12^ vg / mL; Addgene) and TG-controls received sterile saline.

### Experiment 1 – Effect of chronic VTA DA neuron stimulation on biobehavioural sensitisation

#### Behavioural testing

Female (n_hM3D(Gq)_ = 6; n_control_ = 6) and male (n_hM3D(Gq)_ = 6; n_control_ = 7) rats received CNO (1.0 mg/kg) daily for 39 consecutive days (herein referred to as “stimulated rats”). Locomotor activity was then assessed 7 days later during three, 60-minute baseline sessions, and then in response to cocaine (10.0 mg/kg, i.p.), CNO (5.0 mg/kg, i.p.), and saline (0.9%; 1.0 mL/kg, i.p.) challenges (see Figure 1A). Drugs were administered immediately prior to each test session.

**Figure 1.**
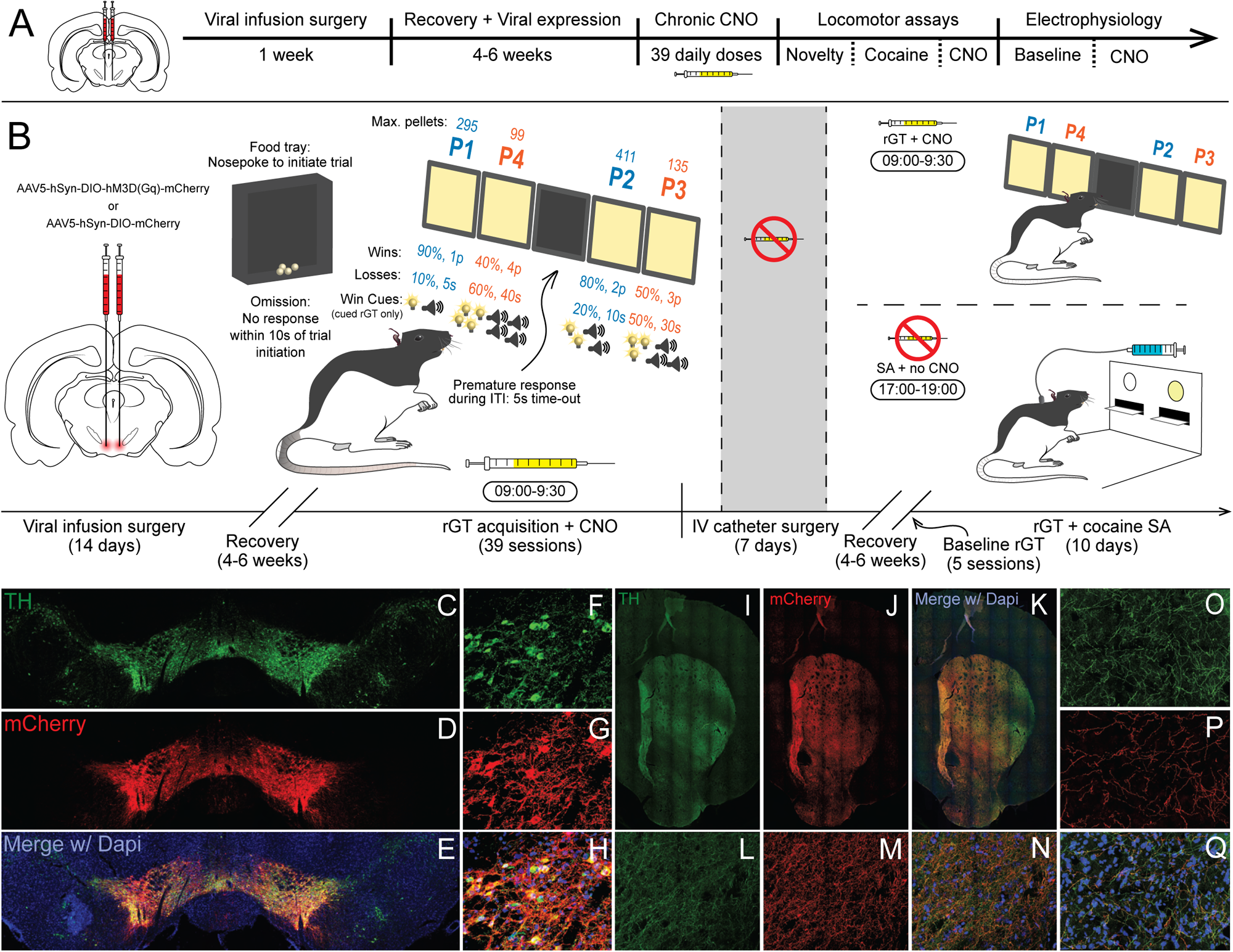
Experimental timeline. **(A)** Experiment 1. The excitatory DREADD AAV5-hSyn-DIO-hM3D(Gq)-mCherry was sterotaxically delivered into the bilateral VTA of female and male TH::Cre rats, which were then given 4–6 weeks to recover with their cage mates with ad libitum food. Following recovery from a stereotaxic surgery, rats were given daily CNO (1.0 mg/kg; i.p.) in their home cages. CNO administration was then ceased for 7 days before locomotor reactivity to novelty, cocaine (10.0 mg/kg; i.p.), and CNO (5.0 mg/kg; i.p.) were assessed. After locomotor testing, rats were surgically prepared for anesthetized electrophysiology and the effect chronic CNO on dopamine neuron physiology was assessed. Responsivity of dopamine neurons to acute CNO were further assessed, to validate the activity of the DREADD. **(B)** Experiment 2. Following recovery from a stereotaxic surgery identical to that of experiment 1, rats were food restricted and began daily rGT training on either the cued or uncued rGT, where CNO (1.0 mg/kg; i.p.) was delivered 30 min prior to the commencement of each session. After 39 rGT sessions, rats were implanted with jugular vein catheters, singly housed, and allowed to recover in their home cages for 1 week. Rats then completed five baseline sessions of rGT with CNO on board, before entering a phase of concurrent cocaine self-administration and rGT training. Each morning, CNO was administered 30 min prior to rGT training. Following each rGT session, rats were returned to their cages for 8 h and, later that evening, placed in different operant boxes and allowed to self-administer intravenous cocaine for 2 h (0.5 mg/kg/infusion; FR1). After 10 consecutive days of concurrent rGT and cocaine self-administration, rats were euthanized, and their brains were processed for immunohistochemistry. **(C-Q)** DREADD expression in VTA dopamine neuron somas and projection regions in the striatum. **(C)** Stitched tilescan of 10X micrographs showing tyrosine hydroxylase (TH)-positive neurons of the VTA in green, **(D)** mCherry-tagged neurons indicating DREADD expression in red, and **(E)** both channels merged with DAPI. **(F-H)** 63X micrographs showing colocalization of mCherry and TH, demonstrating selective transduction of the DREADD in VTA DA neurons. **(I)** Stitched tilescan of 10X micrographs showing TH- and **(J)** mCherry-positive terminals in the striatum, **(K)** with both channels overlaid with DAPI. **(L-N)** 63X micrographs showing colocalization of mCherry and TH in ventral and **(O-Q)** dorsomedial striatum.

#### Electrophysiology

Five days after the last locomotor testing session, rats were implanted with jugular catheters and allowed to recover for 2 days before being prepared for anesthetized electrophysiology (see Figure 1B). A recording electrode was lowered into the VTA to record the electrophysiological properties of spontaneously active dopamine neurons. Once isolated, we recorded from each dopamine neuron for 2 to 3 minutes, sampling (1) the number of spontaneously active dopamine neurons recorded in each track (an index of population activity), (2) firing rate, and (3) the proportion of spikes in bursts (i.e., the occurrence of two spikes with an interspike interval < 80 ms, with its termination defined by an interval > 160 ms). After sampling through all tracks, we chose a previously-identified representative dopamine neuron to record during the CNO challenge. Baseline activity was recorded from this cell for three minutes before administering CNO (1mg/kg, i.p.). Recording continued for a further 15 minutes. See supplemental methods for further details on recording apparatus and parameters.

### Experiment 2 – Effect of sensitization on risk taking before and after cocaine self-administration

#### Behavioural testing

A separate cohort of female (n = 76) and male (n = 58) rats were randomly assigned to training in either the cued or uncued rGT (see supplemental methods and Figure 1B for detailed task descriptions). Rats received one training session per day, 5-7 days per week, for 39 sessions. Thirty minutes prior to each rGT session (which took place from 09:00 – 09:30), rats received IP injections of CNO (1.0 mg/kg in 5% DMSO), thereby matching the CCS regimen of experiment 1. After the final rGT session, a subset of rats (n_female_ = 40; n_male_ = 30) matched for baseline decision making score across experimental group were implanted with jugular vein catheters as per our previously published methods (see supplemental materials for detailed surgical method). After allowing 1 week of recovery, rats resumed daily rGT sessions, with daily CNO still administered 30 minutes prior. After 5 baseline sessions of rGT training, rats began cocaine self-administration (0.5 mg/kg/infusion; FR1) sessions, which took place from 17:00 – 19:00. rGT testing continued as usual the following morning from 09:00 – 09:30. CNO activates DREADDs and modulates behaviour for 40-70 minutes [34, 35] (see Figure 1B). As such, we minimised the chances that CNO would be active during the cocaine self-administration sessions by delivering CNO 8.5 hours earlier. After 10 consecutive days containing both cocaine self-administration and cued rGT sessions, rats were euthanized and the brains processed for immunohistochemistry (see supplemental materials and Figure 1C-Q).

#### Statistical analysis

See supplemental methods.

## Results

### Experiment 1 – Effect of chronic VTA DA neuron stimulation on biobehavioural sensitisation

#### Novelty-, cocaine-, and CNO-evoked locomotion

Chronic chemogenetic stimulation of VTA DA neurons increased locomotor activity in the first, but not subsequent baseline sessions (Figures 2A & 2C; group x BL session: F_2,48_ = 8.99, p < 0.01; BL1_female_: t_10_ = 3.90, p = 0.01; BL1_male_: t_11_ = 2.124, p = 0.04), suggesting heightened locomotor responsivity to novelty. Experimental rats also showed a stronger locomotor response to cocaine (Figures 2B & 2C: drug x group x sex: F_2,72_ = 3.54, p = 0.03; COC [control vs. hM3D(Gq)]-males: t_11_ = 2.52, p = 0.02; -females: t_10_ = 8.35, p < 0.01). 5 mg/kg CNO i.p. only increased locomotor activity in experimental rats (CNO [control vs. hM3D(Gq)]-males: t_11_ = 4.47, p < 0.01; -females: t_10_ = 4.88, p < 0.01)).

**Figure 2.**
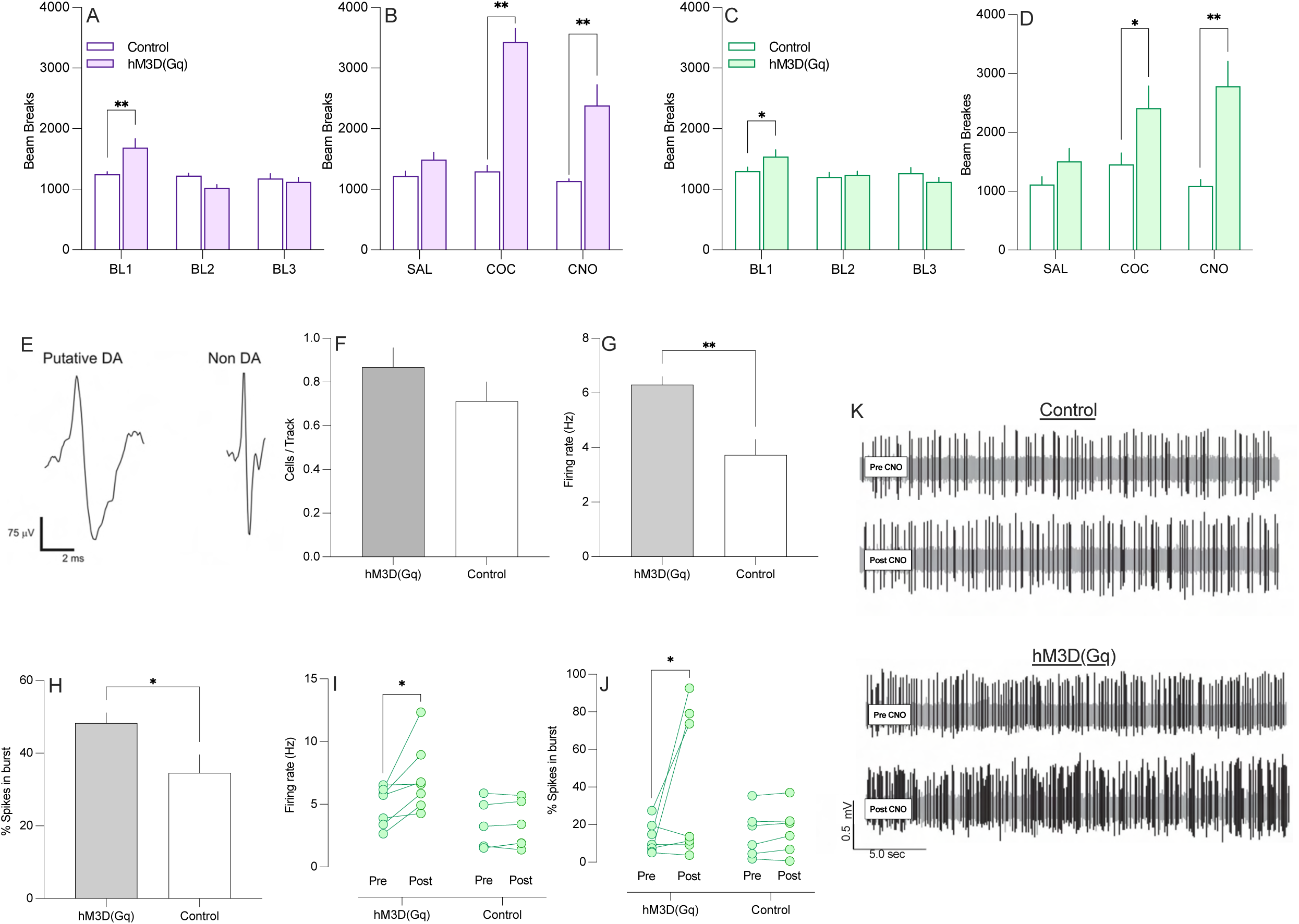
Sensitizing effect of chronic VTA DA neuron stimulation. **(A)** Stimulated females exhibited heightened locomotor in a novel environment [i.e., in the first baseline (BL) session: BL1_female_: t_10_ = 3.90, p = 0.01; BL2 _female_: t_10_ = 1.78, p = 0.23; BL3 _female_: t_10_ = 0.49; p = 0.94and **(B)** showed greater locomotor reactivity than controls to a cocaine (COC) and CNO challenge (stimulated females SAL vs. CNO: t_6_ = 2.70, p = 0.04; controls: t_6_ = 1.77, p = 0.13. **(C)** Stimulated males similarly showed heighted locomotor reactivity to novelty (BL1_male_: t_11_ = 2.124, p = 0.04; BL2_male_: t_11_ = 0.28, p = 0.78; BL3_male_: t_11_ = 1.26; p = 0.22), and **(D)** COC, and CNO challenges (stimulated males SAL vs. CNO: t_6_ = 2.66, p = 0.04; controls: t_7_ = 0.29, p = 0.79). **(E)** Representative waveform for slow firing, long spike width (> 2.5 msec), putative DA neurons (left) and fast-firing, thin spiking (presumably GABAergic) neurons obtained with extracellular recording in the VTA. **(F)** Stimulated rats did not differ from control rats in the number of spontaneously active dopamine neurons detected per recording track (i.e., tonic activity). Stimulated rats showed enhanced markers of phasic activity as evidenced by an increased **(G)** firing rate and **(H)** % of spikes in burst mode. **(I)** In DREADD-expressing rats only an acute CNO challenge increased the firing rate and **(J)** % of spikes in burst mode. **(K)** Representative traces of VTA DA neuron firing in control and stimulated rats at baseline and following administration of CNO. Bars represent group means ± SEM. Pairs of points with connecting lines represent the pre- and post-CNO values of individual animals. Individual point pairs represent individual animals. “*” indicates p < 0.05. “**” indicates p < 0.01.

#### Spontaneous and CNO-evoked dopamine neuron firing

Stimulated and control rats did not differ in the number of spontaneously active dopamine neurons per recording track (Figure 2F; t_15_ = 1.55, p = 0.14). However, dopamine neurons in stimulated rats exhibited a higher firing rate (Figure 2G; t_15_ = 3.85, p < 0.01), and these cells displayed greater percentage of their spikes in burst firing modes (Figure 2H; t_15_ = 2.30, p = 0.04). Rates of mortality for females under chloral hydrate anesthesia were high, therefore Figures 9F-H show males (stimulated: n = 6; control: n = 7) and females (stimulated: n = 2; control: n =2) collapsed together. No female data was collected in the following within-cell analyses.

At the end of the last recording track, individual dopamine neurons were isolated and baseline firing rates recorded. Subsequent treatment with CNO increased the firing rate of dopamine neurons in stimulated rats (pre vs. post: t_7_ = 2.82, p = 0.03) but did not affect this measure in control rats (pre vs. post: t_6_ = 0.31, p = 0.77) (Figure 2I). CNO also increased the percentage of observed spikes in burst mode firing in stimulated rats (pre vs. post: t_7_ = 1.83, p = 0.046), while having no effect in control rats (pre vs. post: t_6_ = 0.157, p = 0.88) (Figure 2J). The null effects of CNO in controls confirmed a lack of off-target effects on dopamine neural activity at this dose. Collectively, these data confirm that repeated chemogenetic stimulation of dopamine neurons lead to long-lasting increases in the excitability of these cells. They further confirm the effectiveness of CNO in activating dopamine neurons expressing Gq DREADD receptors.

### Experiment 2 – Effect of sensitization on risk taking before and after cocaine self-administration

#### Decision making

Consistent with previous reports (for review, see [33]), rats performing the cued rGT had lower scores overall, indicative of greater risky choice (task: F_1,127_ = 20.51, p < 0.01). Chemogenetic stimulation of VTA DA neurons also differentially impacted decision score in males and females (sex x group: F_1,38_ = 1.91, p < 0.01). Likewise, both sex and task condition influenced the effect of VTA DA neuron stimulation on choice of P1-P4 across training (session x choice x group x task x sex: F_114,14934_ = 217.44, p = 0.02). We therefore analyzed decision making score and choice pattern separately for each sex and each task.

### Uncued rGT

#### Females

Stimulation of VTA DA neurons significantly decreased decision making score in females, indicative of greater risky choice (Figure 3A). This was evident in both the first (group_early_: F_1,35_ = 6.42, p = 0.02) and last (group_late_: F_1,35_ = 5.93, p = 0.02) five sessions of training. The rate of change in decision making score did not differ across experimental condition over training (group difference in slopes: F_1,1439_ = 3.29, p = 0.07), with both groups showing stable patterns of overall risk-preference. The lower score in stimulated females was driven partly by a sustained preference for the riskiest option, P4 (Figure 3E; group_early_: F_1,35_ = 6.53, p = 0.02; group_late_: F_1,35_ = 7.32, p = 0.01) and a trend-level increase in choice of the other risky option, P3, over time (Figure 3D; group difference in slopes: F_1,1439_ = 3.19, p = 0.07; slope_control_ = 0; slope_hM3_ = 0.11). In addition, control females developed a relatively greater preference for the best option, P2, with increased training (Figure 3C; group difference in slopes: F_1,1439_ = 13.53, p < 0.01; slope_control_ = 0.82; slope_hM3_ = 0.35), such that choice of this option was significantly lower in the experimental group across the last five sessions (group_late:_ F_1,35_ = 6.49, p = 0.02). Selection of the second most optimal choice, P1, was comparable across groups in all analyses (Figure 3A; group_early_: F_1,35_ = 1.55, p = 0.22; group_late_: F_1,35_ = 2.12, p = 0.15)

**Figure 3.**
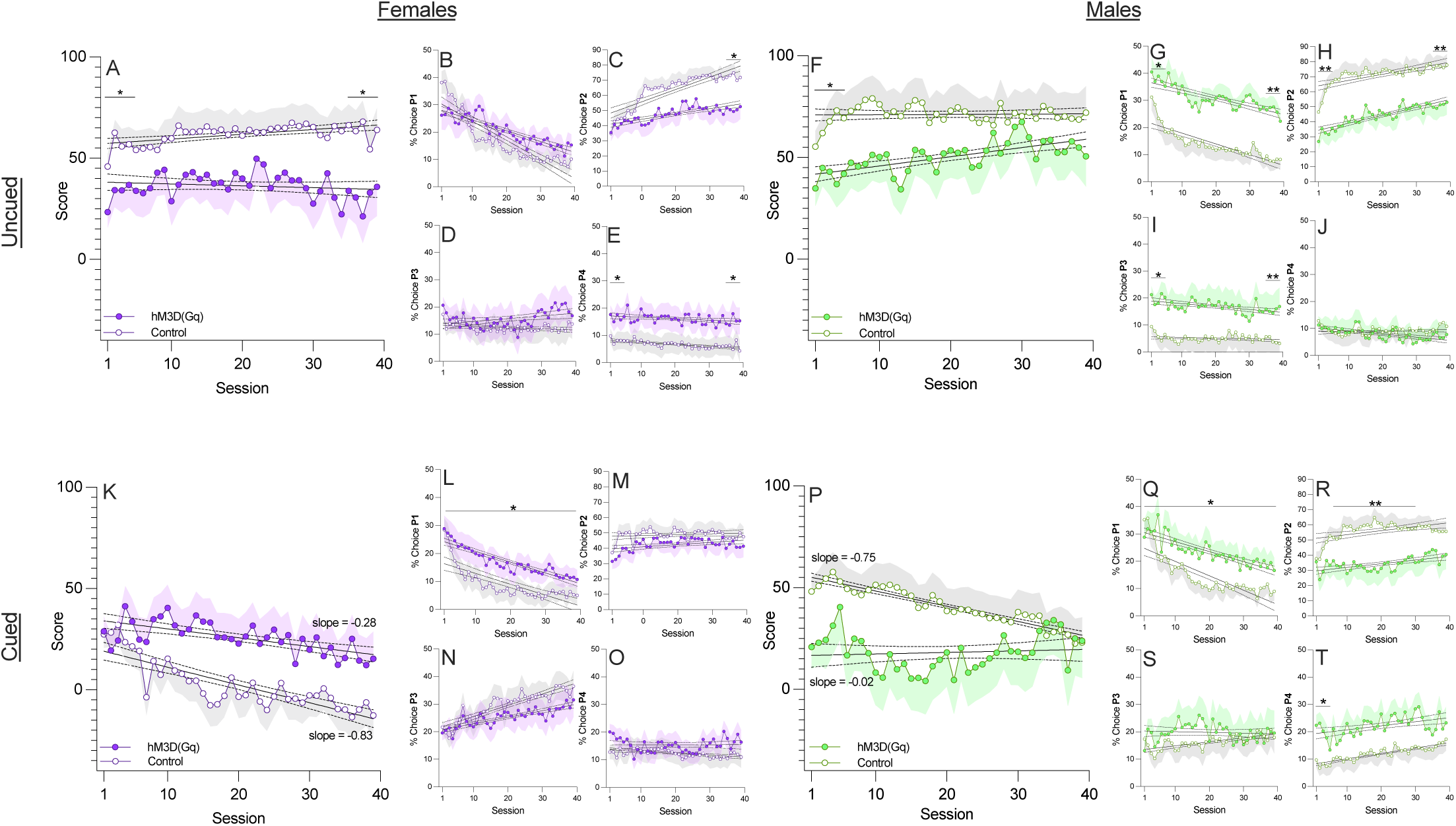
Effect of chronic VTA DA neuron stimulation on decision making. In the uncued rGT, **(A)** stimulated females exhibited a lower decision making score in early and in late training. **(B)** The choice of P1 was not affected by VTA DA neuron stimulation, while **(C)** Stimulated animals showed a decreased preference for P2 in late training. **(D)** Stimulation had no effect on P3 but **(E)** increased the preference for P4 in early and late training. **(F)** Chronic VTA stimulation decreased decision making score in early training, but stimulated males did not differ from controls by late training. **(G)** In early and late training stimulated animals showed enhanced preference for P1, **(H)** reduced preference for P2, and **(I)** increased preference for P3. **(J)** The choice of P4 was unaffected. In the cued rGT **(K)** stimulated females showed a slower rate of decline in decision making score than controls. **(L)** Stimulated females showed an increased preference for P1 throughout training, but did not differ from controls in the choice of **(M)** P2, **(N)** P3, or **(O)** P4. **(P)** Stimulated males exhibited a low decision making score from early training, whereas controls required extensive training to arrive at an equivalently low score. **(Q)** Stimulated males showed an increased preference for P1 throughout training, **(R)** a decreased preference for P2 in mid training, **(S)** no difference in the choice of P3, and **(T)** an increased preference for P4 in early training. Shaded areas represent the SEM. Regression lines are bound by dashed lines denoting the 95% CI. “*” indicates p < 0.05. “**” indicates p < 0.01.

#### Males

While chemogenetic activation of VTA DA neurons also elevated risky choice in males, as indicated by lower decision-making score at the outset of training (Figure 3F; group_early_: F_1,30_ = 5.94, p = 0.02), score increased over time (group difference in slopes: F_1,1244_ = 6.87, p = 0.01; slope_control_ = 0; slope_hM3_ = 0.45) to match that of controls by the final 5 sessions (group_late_: F_1,30_ = 1.59, p = 0.22). The initial reduction in score seen in experimental animals can be attributed to more frequent choice of P3 (Figure 3I; group_early_: F_1,30_ = 22.07, p < 0.01) combined with lower choice of P2 (Figure 3H; group_early_: F_1,30_ = 10.84, p < 0.01; group_late_: F_1,30_ = 7.71; p < 0.01). Both these effects persisted throughout training (group_late_ – P3: F_1,30_ = 5.75, p = 0.02; P2: F_1,30_ = 7.71, p = 0.01), although choice of P2 increased over time in both groups (Figure 3H; group difference in slopes: F_1,1244_ = 0.85, p = 0.36; slope_control_ = 0.40; slope_hM3_ = 0.53). Choice of P1 was also consistently higher in the stimulated group (Figure 3G; group_early_: F_1,30_ = 4.27, p = 0.05; group_late_: F_1,30_ = 7.91, p = 0.01). The progressive increase in score in experimental animals is likely driven by a gradual increase in preference for P2, relatively robust choice of P1, and a relative decrease in preference for P4 over time (group difference in slopes: group_slope_: F_1,1244_ = 5.01, p = 0.03; slope_control_ = 0.01; slope_hM3_ = -0.12), which drove the null effect in score in late training.

#### Cued rGT – females

Decision-making score was initially comparable across groups (Figure 3K; group_early_: F_1,38_ = 0.03, p = 0.87), yet decision making score declined 3 times more rapidly in the control group (Figure 3K; group difference in slopes: F_1,1553_ = 3.903, p = 0.048; slope_control_ = - 0.83; slope_hM3_ = -0.28). However, this did not result in a significant difference between the groups in the last five sessions of training (Figure 3K; group_late_: F_1,38_ = 2.32, p = 0.14). At no point did stimulated females significantly differ from controls in the choice of P2, P3, or P4 (group: all F < 0.251, all p > 0.62), yet they showed a greater preference for P1 (group: F_1,38_ = 4.79, p = 0.04) over the entire course of training. As such, potentiating VTA DA neuron activity in females may slow the rate at which reward-concurrent cues drive risky choice.

#### Cued rGT – males

Initially, decision making score was lower in the experimental group (Figure 3P; group_early_: F_1,28_ = 4.20, p = 0.05) in early training. With continued training, the score of control males declined 25 times more rapidly (Figure 3P; group difference in slopes: F_1,1165_ = 8.12, p < 0.0; slope_control_ = -0.75; slope_hM3_ = -0.02) and matched that of stimulated males by the last 5 sessions (Figure 3P; group_late_: F_1,28_ = 0.01, p = 0.93). The initial difference in score was likely driven by a stronger preference for P4 in the experimental group that was no longer evident by the end of training (Figure 3T; group_early_: F_1,28_ = 8.14, p = 0.01; group_late_: F_1,28_ = 0.88, p = 0.36). Though we did not plan the comparison, visual inspection suggested that stimulated males chose P2 less than controls during mid-stage training; a post-hoc analysis of sessions 6-30 confirmed this observation (Figure 3R; group: F_1,28_ = 7.76, p = 0.01). Stimulated males also chose P1 more frequently (F_1,28_ = 9.22, p = 0.01)

#### Impulsivity

Omnibus ANOVA revealed a session x sex x task x group interaction (F_38,4826_ = 1.57, p = 0.01), prompting us to analyze the effect of group in each task and sex separately. In both task variants, stimulated females consistently made more premature responses after the first five sessions (Figure 4A, 4C: uncued: group_mid_: F_1,35_= 8.70, p = 0.01, group_late:_ F_1,35_= 7.75, p = 0.01; cued: group_mid_: F_1,38_ = 12.94, p < 0.01, group_late:_ F_1,38_= 6.14, p < 0.02), while stimulated males did not differ from controls at any stage (Figure 4B, 4D: all F’s ≤3.04, all p’s ≥ 0.09)

**Figure 4.**
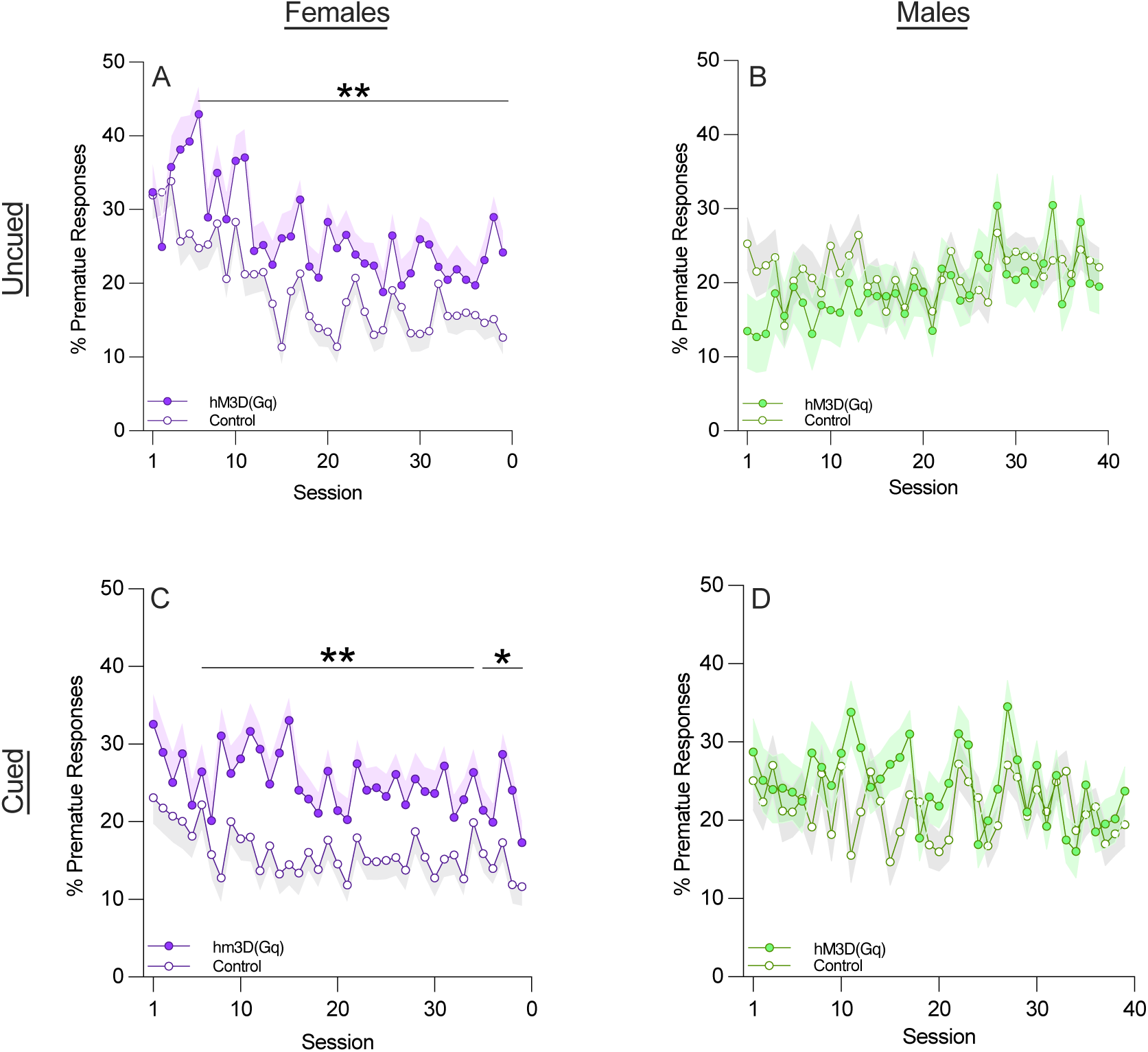
Chronic VTA DA neuron stimulation increases motor impulsivity in females. In the uncued rGT, **(A)** stimulated females make a greater proportion of premature responses than controls throughout mid and late training, while **(B)** stimulated males do not differ from controls. In the cued rGT **(C)** stimulated females also make a greater proportion of premature responses than controls throughout mid and late training, **(D)** while males are not affected. Shaded areas represent the SEM. “*” indicates p < 0.05. “**” indicates p < 0.01.

#### Other Variables

See supplemental results.

#### Cocaine self-administration

As per our previous report [31], training in the cued rGT enhanced the self-administration of cocaine (see Supplemental Figure 2I). Independent of sex (sex: F_1,66_ = 0.01, p < 0.94), stimulation affected cocaine self-administration differentially between tasks (group x task: F_1,66_ = 7.37, p = 0.01). In the cued rGT, stimulated rats took significantly more infusions (Figure 5B; group: F_1,42_ = 24.08, p < 0.01) and made more responses for cocaine than controls (Figure 5D; group: F_1,42_ = 22.84, p < 0.01). Stimulated animals did not differ from controls in the uncued task (Figures 5A & 5C; infusions - group: F_1,28_ = 3.62, p = 0.07; active responses - group: F_1,28_ = 2.78, p = 0.11). In both the cued (F_1,42_ = 4.51, p < 0.04) and uncued (group: F_1,28_ = 11.19, p < 0.01) rGT, stimulated animals made more inactive responses than controls (Figures FC & 5D). To control for this, we conducted an additional analysis comparing the difference of active and inactive responses across groups, which reaffirmed the group-wise differences we originally detected (group: cued rGT: F_1,42_ = 11.93, p < 0.01; -uncued rGT: F_1,28_ = 2.09, p = 0.16). See supplemental results for data analyzed by sex.

**Figure 5.**
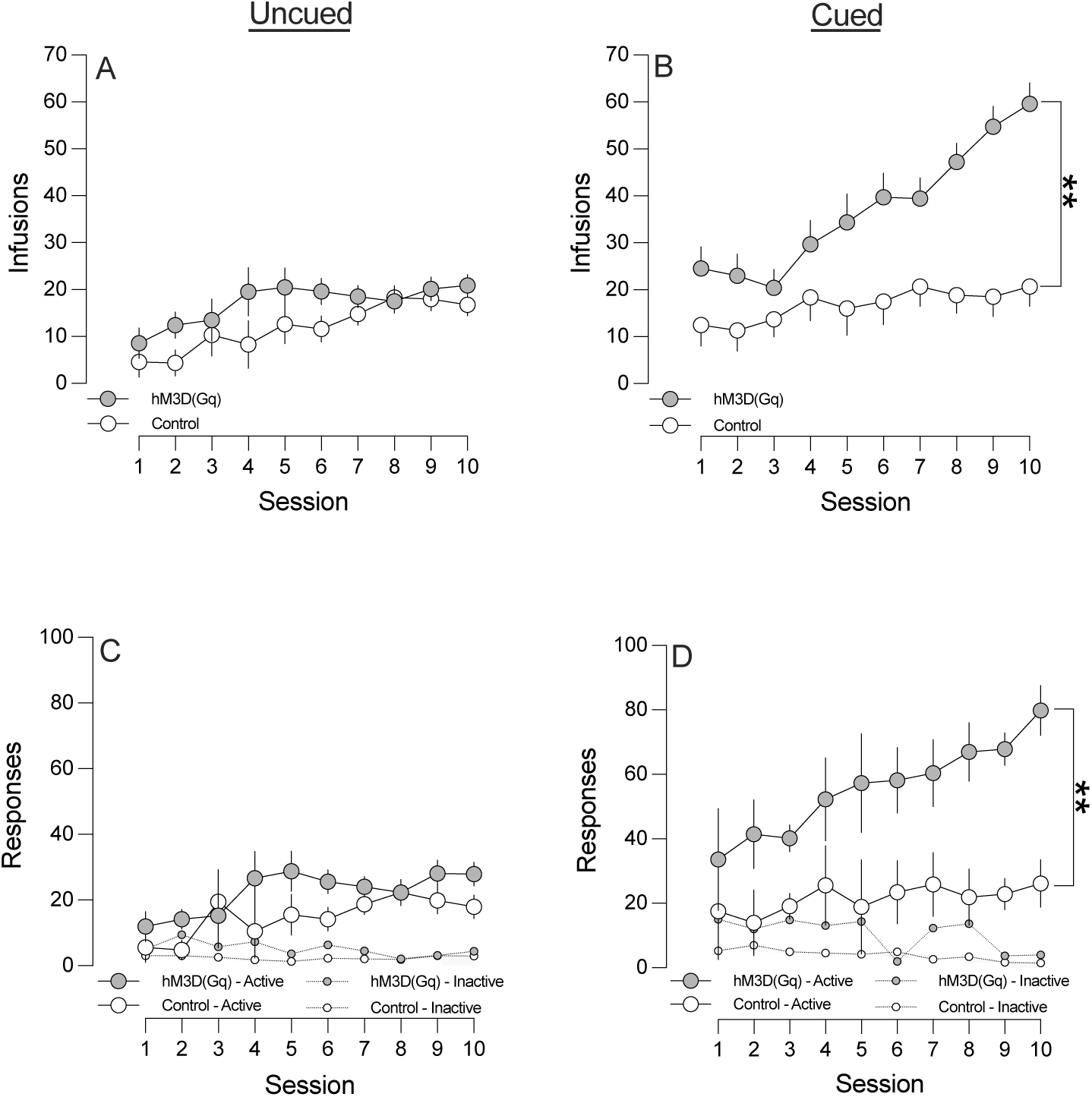
Chronic VTA DA neuron stimulation potentiates cocaine self-administration in the cued rGT. **(A)** Stimulation had no effect no effect on the number of infusions of cocaine taken by animals trained in the uncued rGT. **(B)** Stimulated animals trained in the cued rGT took more infusions of cocaine than control counterparts. **(C)** Stimulated animals trained in the uncued rGT did not differ from controls in the number of responses made upon the active or inactive lever. **(D)** Stimulated animals trained in the cued rGT made more responses upon the active lever but did not differ from controls in the number of inactive responses made. Error bars represent the SEM. “**” indicates p < 0.01.

#### Decision making during cocaine self-administration

Cocaine self-administration differentially affected decision making across group and task, but we detected no sex differences (score – session x group x task: F_14,994_ = 1.72, p = 0.05; choice profile – session x choice x group x task: F_42,2856_ = 1.43, p = 0.04). In the uncued rGT, cocaine self-administration did not affect decision making score (Figure 6A; session: F_14,420_ = 0.69, p = 0.78; session x group: F_14,420_ = 1.06, p = 0.39). Although cocaine self-administration decreased the preference for P4 in stimulated rats (session x group: P_14,420_ = 3.22, p < 0.01; pre vs. post_ctrl_: t_15_ = -1.59, p = 0.07; pre vs. post_hM3_: t_15_ = 2.22, p = 0.02), no other choice was significantly affected in either group (P1-P3, session, session x group: all F’s ≤ 1.68, all p’s ≥ 0.14)

**Figure 6.**
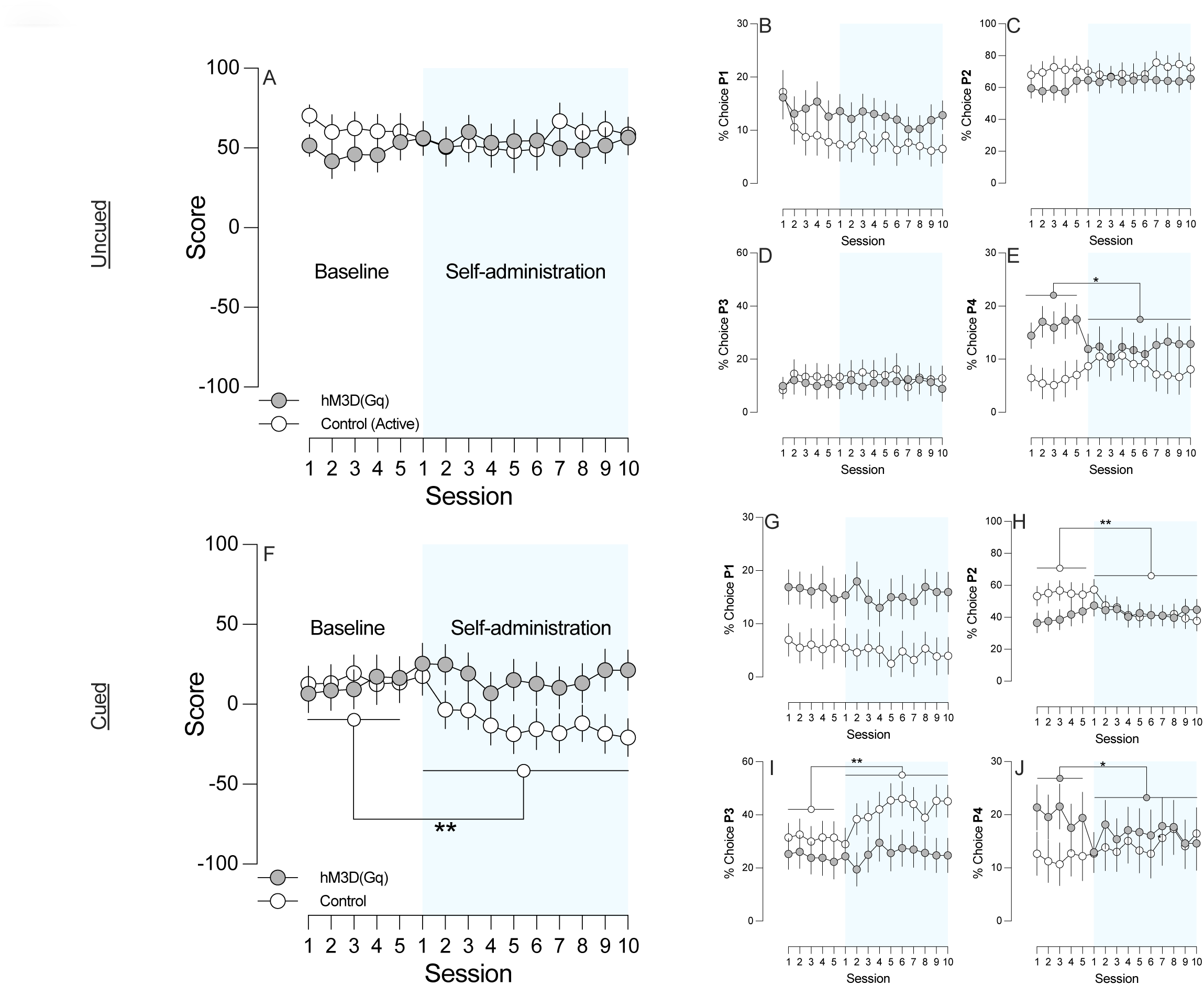
Chronic VTA DA neuron stimulation modulates the effect of cocaine on decision making. **(A)** In the uncued rGT, neither controls nor stimulated animals displayed cocaine-induced changes in decision making score, **(B)** the choice of P1, **(C)** P2, **(D)** or P3. **(E)** In stimulated animals, cocaine self-administration decreased the preference of P4. **(F)** In the cued rGT, cocaine self-administration decreased decision making score in control animals **(G)** The choice of P1 was unaffected. **(H)** Control animals showed a reduction in the preference for P2 **(I)** and an increase in the preference for P3. **(J)** Stimulated animals showed a decrease in the preference for P4. Error bars represent the SEM. “*” indicates p < 0.05. “**” indicates p < 0.01.

In the cued rGT, decision making score decreased in control rats following cocaine self-administration, but not in stimulated rats (Figure 6F; group x session: F_14,630_ = 5.47, p < 0.01; pre vs. post_ctrl_: t_24_ = 3.46, p < 0.01; pre vs. post_hM3_: t_21_ = -1.65, p = 0.12); this effect was driven by decreased choice of P2 (Figure 6H; session x group: F_14,630_ = 5.06, p < 0.01; pre vs. post_ctrl_: t_24_ = 3.91, p < 0.01; pre vs. post_hM3_: t_21_ = -0.51, p = 0.31) and increased choice of P3 (Figure 6I; session x group: F_14,630_ = 2.81, p < 0.01; pre vs. post_ctrl_: t_24_ = -3.83, p < 0.01; pre vs. post_hM3_: t_21_ = -1.19, p = 0.12). In stimulated rats, cocaine self-administration again decreased preference for P4 without affecting overall score (Figure 6J; session x group: F_14,620_ = 2.22, p = 0.01; pre vs. post_ctrl_: t_24_ = -1.48, p < 0.15; pre vs. post_hM3_: t_21_ = 2.43, p = 0.02).

#### Impact of cocaine-self administration on other variables

See supplemental results.

## Discussion

The present set of experiments demonstrate the critical impact of a sensitized dopamine system on decision making, cocaine self-administration, and the interaction thereof. We show that a regimen of repeated, daily stimulation of VTA DA neurons is sufficient to sensitize the locomotor response to both cocaine and CNO, and elevated phasic but not tonic activity. By chemogenetically sensitizing the DA system, we caused behaviour in the uncued rGT to mimic the cue-induced risk taking we report here and elsewhere in the cued rGT [12]. However, unlike repeated engagement with the cued rGT, chemogenetic sensitization of VTA DA neurons was not sufficient to enhance cocaine self-administration following experience with the uncued rGT. Nevertheless, combining these two manipulations significantly potentiated cocaine-taking. Males in the experimental group also reached an asymptotic level of risk preference more rapidly on the cued rGT, further indicating synergy between DA neuron stimulation and cue-driven risky choice. In contrast, the opposite effect was observed in females, such that dopaminergic sensitisation appeared to counter the risk-promoting effect of the cues. In females, the addition of reward paired cues to the rGT therefore dramatically alters the role played by the dopamine system in establishing choice preferences. Contrary to our predictions, VTA DA sensitization prevented rather than enhanced the progressive increase in risky decision making typically observed on the cued rGT the morning after cocaine self-administration sessions. Together, these findings may help us understand how ingestion of psychostimulants and engagement with heavily cued gambling games interact to modulate dopaminergic signaling, leading to a degree of substitutability and cross-dependence.

### Chemogenetic stimulation sensitizes the mesolimbic dopamine system

To verify that our DREADD-based manipulation resulted in a behavioural phenotype comparable to the classic demonstrations of psychomotor sensitization [15, 36], we confirmed that repeated chemogenetic stimulation of VTA dopamine neurons sensitized the psychomotor stimulant effects of cocaine. This observation closely mimics the effects of chronic electrical stimulation of the VTA [37]. CNO acutely promoted locomotion selectively in DREADD-expressing rats, similar to previous reports of pharmacological [38], electrical [39], optical [40], and chemogenetic [41] activation of the dopamine system. Consistent with the null effect of Gq stimulation on baseline locomotion, dopamine neuron population activity (i.e., cells/track, an index of tonic dopamine activity) was similar between DREADD-expressing and control rats. However, markers of phasic activity (i.e., firing rate and cells in burst mode) were elevated. We also observed CNO-evoked increases in measures of phasic dopamine activity, further validating use of the DREADD as a manipulation which enhances intrinsic excitability. Greater phasic dopamine activity has previously been observed in rats that were more hyperactive in a novel environment and self-administered more cocaine [42]. Likewise, chemogenetic enhancement of phasic dopamine activity also led to enhanced locomotor response to novelty and greater cocaine self-administration in the present study, indicative of a pro-addiction phenotype.

### Divergent dopaminergic control of decision making in females and males: critical cue effect

In the absence of reward cues, stimulating VTA DA neurons promoted risk-taking in both sexes, consistent with reports that artificially driving the dopamine system biases males [43, 44] and females [45] toward risk taking in probabilistic decision making tasks. However, when reward-cues were present, VTA DA neuron stimulation facilitated the development of a risk-preferring strategy in males while attenuating it in females. The opposing direction of these effects mirrors chemogenetic inhibition of VTA DA neurons during acquisition of the cued rGT [32], as this manipulation attenuates risk-preference in males and exacerbates it in females. Because reward predictions errors are similarly encoded in VTA DA neurons of females and males [46, 47] and both sexes show identical dopaminergic responses to reward cues [48], females must use this teaching signal to guide choice very differently than males. Dramatic sex differences in the terminal field structures of VTA DA projections (e.g., the basolateral amydgala [49]) may be the key to understanding these differences.

### Sex differences in the dopaminergic modulation of impulsivity

Sensitizing VTA DA neurons also increased impulsive, premature responding in female, but not male, rats in both tasks. Conversely, chemogenetic inhibition of this neuronal population decreased such impulsive responding in males but not females [32]. The critical role of dopamine in regulating motor impulsivity is well-characterised [50–52]. In *male* rats performing the 5-CSRTT, systemic amphetamine potentiates premature responding; this effect is most likely mediated by dopamine in the NAc, as intra-accumbal infusions of amphetamine provoke similar increases in premature responding that are blocked by dopamine but not noradrenaline antagonists [53]. Optogenetic stimulation of the VTA-NAc projection also increases premature responding, while stimulation of the VTA-PFC projection had no effect on impulsivity [54]. However, the behavioural response to dopamine receptor stimulation may be influenced by sex, with females showing a more pronounced increase in premature responding following amphetamine, compared to males [55]. Females may therefore be more susceptible to the “impulsogenic” effects of increased dopaminergic activity, whereas decreases may reduce impulsive responding more prominently in males. Indeed, female rats show lower levels of baseline dopamine efflux in the accumbens and their mesoaccumbal projection is less dense than males [56, 57], aligning with the lower impulsivity observed in female humans and rats alike [58].

### Reward cues increase the influence of sensitization on cocaine self-administration

Rats trained in the cued rGT self-administered more cocaine than their uncued counterparts, and this was potentiated by chronic chemogenetic stimulation of VTA DA neurons. This is in keeping with the finding that rats pre-exposed to an experimenter-administered sensitizing regimen of cocaine go on to self-administer more cocaine [59]. Similarly, more novel or salient settings can exaggerate the behavioral expression of psychomotor sensitization to stimulant drugs [60]. However, if cue-biased risky choice and chemogenetic sensitization of DA neurons work synergistically to promote cocaine-taking, why did cocaine self-administration and chemogenetic sensitization of DA neurons not promote risky decision-making?

We know from cocaine self-administration studies that animals actively titrate the level of “brain cocaine” to optimal levels [61–63]; the same is also true for brain dopamine levels (electrical self-stimulation: [64]; optical self-stimulation: [65]). Therefore, in addition to considering dopaminergic stimulation as a modulator of decision making, it may also be a behavioural goal, the desired level of which is labile (see [66]). As the dopamine system is repeatedly stimulated, either through dopamine neuron sensitisation, or exposure to heavily-cued probabilistic rewards, the preferred level of brain dopamine may increase. When stimulation ceases, the return to a lower, basal dopamine level may become increasingly aversive [67], leading to greater cocaine self-administration. Drug ingestion may not have driven further risk-taking in stimulated rats here because DREADD activation on-task was sufficient to compensate for this negative state. Choice of P4 may have then dropped in these rats because dips in dopamine would be expected to accompany the frequent losses. Had we stopped CNO administration prior to these sessions, we may have seen the predicted escalation of risky choice, as animals would have been forced to boost brain dopamine levels through riskier decision-making.

In sum, these findings demonstrate synergy between engagement with heavily cued gambling games and cocaine-taking at the level of the dopamine system that suggest pathways to cross-dependence. This study also highlights key differences in how females and males respond to changes in dopamine signalling, which may eventually help us better understand the divergent trajectories of addiction disorders in men and women [68].

## SUPPLEMENTAL MATERIALS

### Supplemental methods

#### Animals

Rats were weaned at PND 21 and were pair- or trio-housed in a climate-controlled colony room on a reverse 12-hour light-dark cycle (lights off 08:00; temperature 21°C). Prior to being assigned to experimental groups, females and males were genotyped to confirm the expression Cre-recombinase. Ear notches were obtained for PCR analyses. To extract DNA, ear notches were lysed using a buffer solution (50 mM pH 8.0 Tris, 2 mM NaCl, 10 mM EDTA, 1% SDS) and Proteinase K (Invitrogen, part number 25530-015). Extracted DNA was stored at - 20°C until PCR was performed. All reactions were performed in 200 µl thin-walled PCR tubes with FASTSTART 28 TAQ DNA POL. DNTPACK reaction mix (Roche, cat #4738357001), primers for Cre (forward: 5’-AGA GTA CAC TGT GGG CAG GA-3’; reverse: 5’-GCA AAC GGA CAG AAG CAT TT-3’) or TH (forward: 5’-CGC TTA CCC CGG AAG AAC AA-3’; reverse: 5’-CCA GCA GAG GTA ATG GAA GAG A-3’). Samples were then placed in a thermocycler and underwent standard cycling protocols (95°C 5 mins; then cycled 35 times: 94°C 30 sec, 63°C 30 sec, 72°C 1 min; holding at 72°C for 10 min; infinite holding at 4°C) and subsequently run on an agarose gel at 90V for 40 min to verify genotypic expression.

#### Behavioural apparatus

Locomotor activity was assayed in 8 operant boxes (30.5 × 24 × 21 cm; MED Associates, St Albans, VT) equipped with four infrared photobeams mounted on the side of each chamber and one photobeam located inside the food receptacle. Each chamber was enclosed within a ventilated sound-attenuating cabinet (Med Associates Inc, VT). The operant boxes were equipped with a fan to provide ventilation and to mask extraneous noise. Locomotor activity was indexed by the number of photobeam breaks that occurred during a session. In a separate laboratory space in the same facility, the rat gambling task and cocaine self-administration behavioural assays were each conducted in 16 separate banks of operant conditioning chambers (30.5 × 24 × 21 cm; Med Associates, St. Albans, VT, USA), located in separate rooms. These operant boxes were housed in similar enclosures to those used to conduct locomotor resting. Set in the curved wall of each box was an array of five nose-poke holes. Each nose-poke unit was equipped with an infrared detector and a yellow light-emitting diode stimulus light. Sucrose pellets (45 mg, Formula P; Bio-Serv) could be delivered at the opposite wall via a dispenser. The cocaine self-administration chambers were identical to the cued rGT chambers apart from an additional cantilevered drug delivery arm and vascular access tether system plus externally mounted variable rate syringe pump (Med Associates Inc., VT). Online control of the apparatus and data collection was performed using code written by CAW or SBF in MEDPC (Med Associates) running on standard IBM-compatible computers.

#### Electrophysiology apparatus

Under chloral hydrate (400 mg / kg, i.v.) anesthesia, a craniotomy was performed in the skull above the VTA, and a recording electrode (2.0 mm OD, ∼1 µm tip diameter, impedance 5-10 MΩ) was lowered into the VTA to record the electrophysiological properties of spontaneously active dopamine neurons. We passed the electrode through 6-9 vertical sampling tracks, identifying putative dopamine neurons based on waveform properties (long-duration (>2.5 ms) bi-or triphasic action potential, often with a break between initial and somato-dendritic spike segments; [69]). The electrode signal was amplified and filtered (500-2000 Hz) using an X-Cell3+ microelectrode amplifier (Fredric Haur Co., Bowdin, ME). Action potential data were acquired, discriminated from noise, stored, and analyzed using Spike 2 software (CED, Cambridge, UK) running on an Intel-based personal computer with a data acquisition board interface (micro 1401 mk II, CED).

#### Jugular catheter surgeries

For experiments 1 and 2, rats were implanted with an intravenous catheter to enable experimenter-administered drugs or drug self-administration, respectively. Specifically, we aseptically implanted catheters constructed of Silastic silicone tubing (Dow Corning via VWR International, Edmonton, AB, Canada), attached to back-mounted cannulae (Plastics One, Roanoke, VA, USA), into the right jugular vein as per our previous published methods [31]. We passed the catheters through the skin subcutaneously and externalized the cannulae between the scapulae. Following surgery, the catheter was filled with a heparin glycerol lock solution (SAI Infusion Technologies, Illinois, USA) and the animals were left to recover for 5-7 days, after which cued rGT testing resumed and cocaine self-administration began. The day prior to the first cocaine self-administration session and immediately after the last self-administration session, catheter patency was tested by delivering 0.1 mL of 10% ketamine HCL (Medisca Pharmaceuticals, British Columbia, Canada). All rats exhibited an immediate loss of muscle tone and sedation, resulting in no rats being excluded on the criterion of catheter patency.

#### Uncued and cued rat gambling task (rGT)

Animals were initially habituated to the operant chambers over the course of two 30-minute exposures, during which sucrose pellets were placed in each of the apertures and animals were allowed to explore the apparatus. Animals were then trained on a variant of the five-choice serial-reaction time task (5-CSRTT) in which one of the five nose-poke apertures was illuminated for 10 s and a nose-poke response was rewarded with a single sucrose pellet delivered to the food magazine. The aperture in which the stimulus light was illuminated varied across trials. Each session consisted of 100 trials and lasted 30 min. Animals were trained on this task until responding reached 80% accuracy and 20% omissions. Once this training was complete, rats then performed a forced-choice variant of the rGT. This training procedure ensured equal exposure to the different reinforcement contingencies associated with each aperture. During each 30-minute cued rGT session, rats sampled between four response holes, each of which was associated with distinct magnitudes and probabilities of sucrose pellet rewards or time-out punishments. The optimal approach in the rGT is to favour options which deliver smaller per-trial gains but lower time-out penalties; consistent choice of the smaller reward options was advantageous due to more frequent rewards, but also less frequent and shorter time-outs, with the two-pellet choice (P2) resulting in the most reward earned per unit time. As detailed in our previous publications, most rats adopt this optimal strategy in the absence of reward-concurrent audiovisual cues [70]. In the cued version of the rGT, a 2-s audiovisual cue was presented concurrent with reward delivery on each option. The cue increased in complexity with the size of the reward: P1 win: P1 hole flashes for 2s at 2.5 Hz, monotone; P2 win: P2 hole flashes for 2s at 2.5 Hz, tone changes pitch once after 1s; P3 win: P3 hole flashes at 5 Hz for 1s, followed by flashing of the two neighboring holes in one of two patterns chosen at random, traylight flashes concurrently at 0.2 Hz for 2s, three different tones used, changing pitch every 0.1 s, in one of two patterns chosen at random; P4 win: P4 hole flashes at 5 Hz for 1s, followed by flashing of all five holes in one of four patterns at random, traylight flashes concurrently at 0.2 Hz for 2s, six different tones used, changing pitch every 0.1 s, in one of four patterns chosen at random. These cues are omitted in the uncued version of the rGT.

Responses made during the ITI were recorded as premature responses, a measure of impulsive action, which resulted in the illumination of the house light and a 5 s time-out penalty after which a new trial could be initiated. If a response was not made into one of the 4 holes during the 10 s stimulus presentation, the trial would be registered as an omission, after which point another trial would begin.

#### Cocaine self-administration paradigm

Animals were trained to lever press for cocaine hydrochloride (0.30 mg/kg/infusion, dose calculated as the salt and dissolved in sterile 0.9% saline; Medisca Pharmaceuticals, British Columbia, Canada) over 10 daily 2-hr sessions. At the start of each self-administration session, two free infusions of solution were given to fill catheters and indicate drug was available. Rats were presented with two levers, one active and one inactive, with an illuminated cue-light situated over the active lever. Using a fixed ratio schedule, responses on the active lever would result in a single 4.5 s infusion in concert with the cue light flashing (50 Hz) and a novel 20 kHz tone (this same audiovisual stimulus was not used in the cued rGT). Following the infusion, animals underwent a 10 s time-out during which drug would not be delivered, the cue light and tone were not presented, but levers would remain extended, and responses monitored. Inactive lever presses, while monitored, had no programmed consequences.

#### Immunohistochemistry

To visualize DREADD expression in VTA dopamine neurons and their striatal projections, we double-labelled 35um sections coronal sections with primary antibodies against mCherry (Cat#ab205402; Abcam; Toronto, ON, Canada; 1:500 for 48 hours) and tyrosine hydroxylase (Cat#AB152; Millipore Sigma; Oakville, ON, Canada; 1:500 for 24 hours) for 24 hours. Sections were then washed in PBS and incubated with secondary antibodies conjugated to Alexa Fluor ® 488 (Cat#A-21103) and Alexa Fluor ® 633 (Cat#A-11034) (Thermo Fischer Scientific; Burnaby, BC, Canada; 1:1000 for both). Sections were then cover-slipped under HARLECO^®^ Krystalon™ mounting medium (Thermo Fischer Scientific; Burnaby, BC, Canada) and visualized using a SP8 WLL confocal microscope (Leica Microsystems, Germany).

#### Statistical analysis

For locomotor analyses, omnibus univariate ANOVAs with sex, group, and baseline session / drug as between-subjects factors (IBM SPSS Statistics Version 28) were first conducted to justify follow-up independent samples t-tests (Prism 9). For electrophysiological data, independent samples t-tests were used to compare between groups, and paired samples t-tests were used to examine pre- and post-CNO DA neuron activity (Prism 9). For all rGT and cocaine self-administration analyses, we followed up significant omnibus outcomes (between-subjects: task, group, and sex; within-subjects: session) with subsequent repeated measures ANOVAs. For decision making, we looked at early (first 5 sessions), late (last 5 sessions), and in certain post-hoc instances, middle (sessions 6-34) training epochs (IBM SPSS Statistics Version 28). We also performed linear regression analyses to compare the change in decision making over the entire course of training (Prism 9). For premature responding and other rGT variables we tested early (first 5 sessions), late (last 5 sessions), and in all instances, middle (sessions 6-34) training epochs. For cocaine self-administration we looked at all 10 sessions in follow-up repeated measures ANOVAs. In the analyses surrounding the impact of cocaine self-administration on rGT performance, significant omnibus ANOVAs were followed up with paired samples t-tests, comparing the means of pre- and post-cocaine epochs.

### Supplemental results

#### Experiment 1: locomotor activity analysis

Compared to saline, 10 mg/kg i.p. cocaine increased locomotor responding in experimental but not control females (drug x group x sex: F_2,72_ = 3.54, p = 0.03; SAL vs. COC-experimental: t_6_ = 11.0, p < 0.01; -control: t_6_ = 1.95, p = 0.10) (Figure 2B). It is possible that pre-treatment with CNO attenuated the locomotor response to cocaine in control females, as others have reported similar findings with cocaine [71] and amphetamine [72]. This dose of cocaine is also lower than is typically used in to evoke sensitization in unmanipulated females [73]. In males, an acute cocaine challenge promoted locomotion in both control (SAL vs. COC: t_7_ = 1.95, = p = 0.05) and stimulated groups (SAL vs. COC: t_6_ = 2.09, p = 0.03) (Figure 2D). However, the locomotor response to cocaine was greater in stimulated males (COC [control vs. hM3D(Gq)]: t_11_ = 2.52, p = 0.02).

#### Experiment 2: Other behavioural variables

##### Trials

In early uncued rGT training, stimulated females and males completed fewer trials (session x sex x group: F_38,4826_ = 1.45, p = 0.036; task: F_1,127_ = 19.48, p < 0.01; females – group_early_: F_1,30_ = 6.78, p = 0.01; males -- group_early_: F_1,35_ = 4.26, p = 0.05) but did not differ from controls in late training (females -- group_late_: F_1,35_ = 1.41, p = 0.24; males – group_late_: F_1,30_ = 2.00, p = 0.17). In the cued rGT, at no point in training did stimulated females differ from controls (group_early_: F_1,35_ = 1.23, p = 0.28; group_late_: F_1,35_ = 1.80, p = 0.19). Males completed fewer trials in early training, but were comparable to controls by late training (group_early_: F_1,27_ = 4.87, p = 0.04; group_late_: F_1,27_ = 0.52, p = 0.48) (Supplemental Figure 1A-D).

#### Omissions

In late uncued rGT training, stimulated females omitted more fewer trials than controls in early training (session x task: F_38,4826_ = 3.56, p < 0.01; task: F_1,127_ = 5.86, p < 0.02; group_late_: F_1,35_ = 7.75, p = 0.01; group_early_: F_1,35_ = 0.38, p = 0.54), while males differed at no point (Supplemental Figure 1F; group_early_: F_1,30_ = 0.99, p = 0.33; group_late_: F_1,30_ = 0.71, p = 0.41). In the cued rGT, stimulated females did not differ from controls (group_early_: F_1,35_ = 3.26, p = 0.06; group_late_: F_1,35_ = 0.07, p = 0.79), whereas stimulated males made more omissions than controls throughout training (group_early_: F_1,27_ = 10.67, p < 0.01; group_late_: F_1,27_ = 3.91, p = 0.05) (Supplemental Figure 1E-H).

#### Choice latencies

In the uncued rGT, stimulated females chose quicker than controls in late training (session x group x sex x task: F_38,4826_ = 1.82, p < 0.01; group_late_: F_1,35_ = 12.04, p < 0.01; group_early_: F_1,35_ = 3.120, p = 0.09). At no point did stimulated males differ from controls (group_early_: F_1,30_ = 1.24, p = 0.27; group_late_: F_1,30_ = 0.12, p = 0.74). In late training, stimulated females chose quicker than controls in the cued rGT (group_early_: F_1,35_ = 2.22, p = 0.14; group_late_: F_1,35_ = 12.59, p < 0.01), whereas stimulated males were slower to make choices throughout training (group_early_: F_1,27_ = 17.22, p < 0.01; group_late_: F_1,27_ = 4.31, p = 0.05) (Supplemental Figure 1I-L).

**Supplemental Figure 1.**
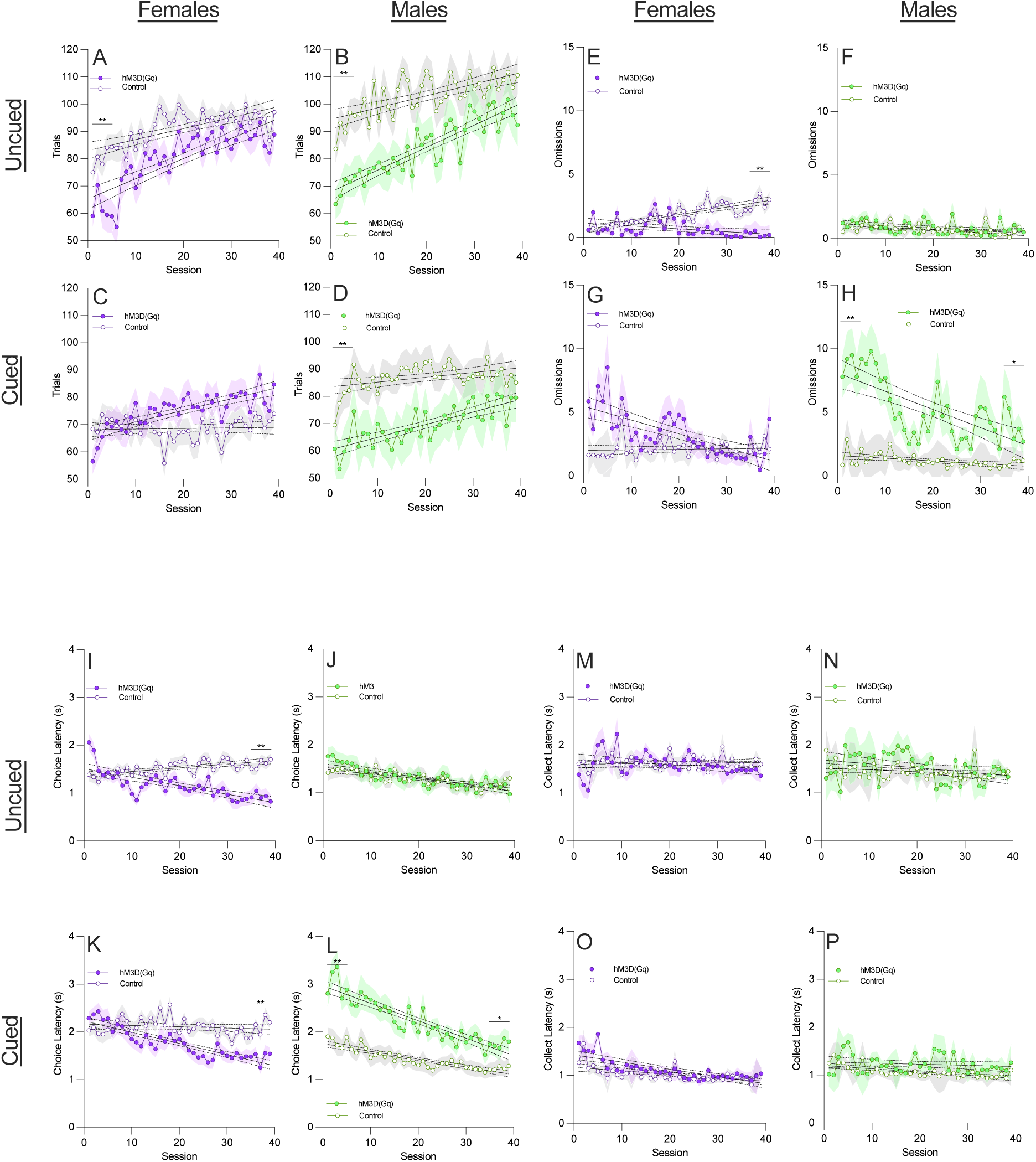
Effect of VTA DA neuron stimulation on auxiliary behavioural measures. **(A)** In the uncued rGT, stimulated females and **(B)** males completed fewer trials than controls in early training but did not differ by late training. **(C)** In the cued rGT, stimulated females did not differ from controls at any point. **(D)** Stimulated males completed fewer trials in early training but did not differ from controls in late training. **(E)** In the uncued rGT, stimulated females omitted fewer trials than controls in late training. **(F)** Stimulated males did not differ from controls at any point in training. **(G)** In the cued rGT, stimulated females did not differ from controls. **(H)** Stimulated males omitted more trials in both early and late training. **(I)** In the uncued rGT, stimulated females were quicker to make a choice than controls by late training, whereas **(J)** males did not differ. **(K)** In the cued rGT stimulated females were quicker to make a choice than controls in late training. **(L)** Stimulated males were slower to choose than controls in early and late training. **(M-N)** In neither the uncued rGT nor **(O-P)** cued rGT did stimulated females or males differ from controls in the speed to collect reward. Shaded areas represent the SEM. “*” indicates p < 0.05. “**” indicates p < 0.01.

**Supplemental Figure 2.**
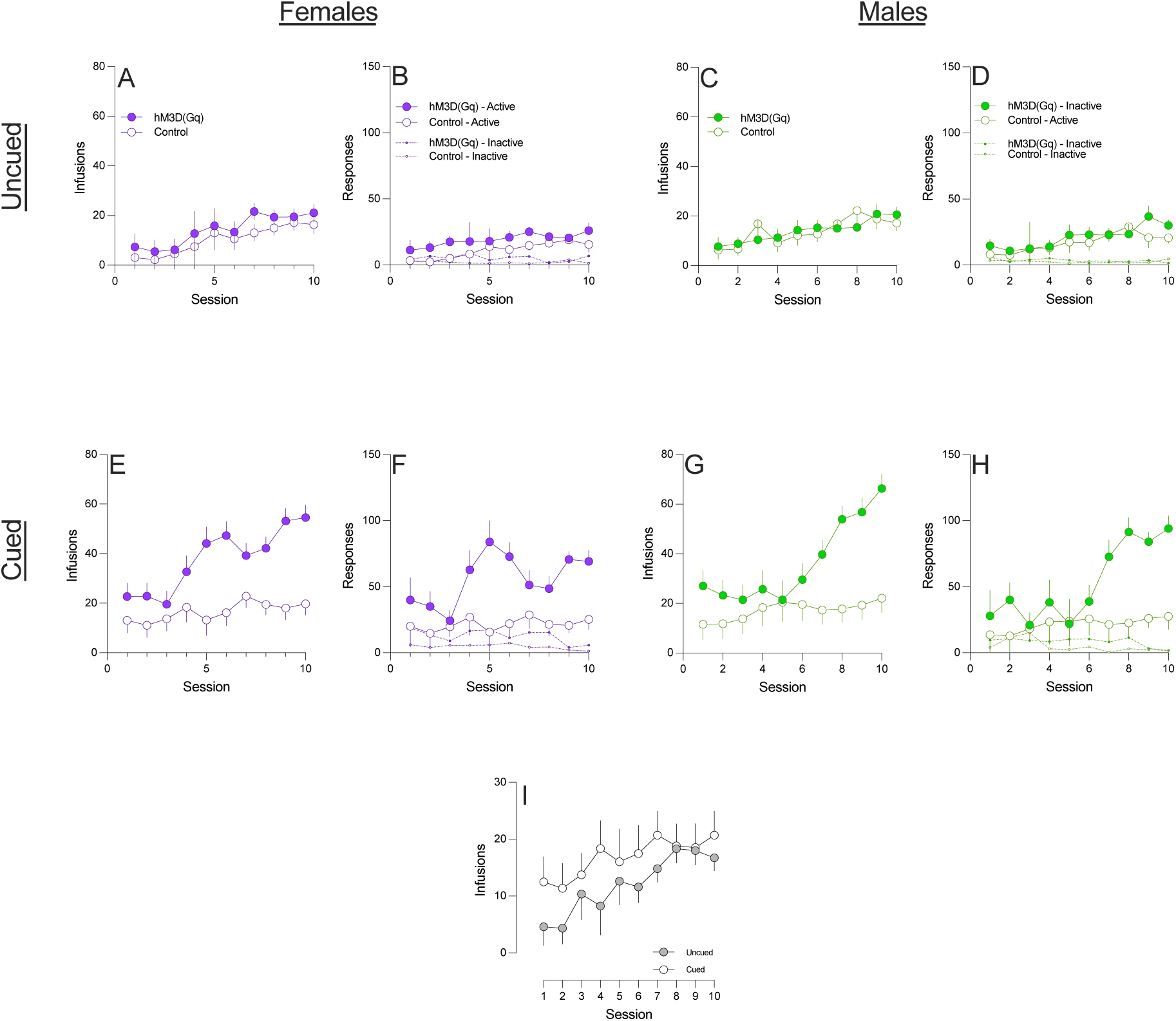
Effect of VTA DA on cocaine self-administration in both sexes. **(A)** In the uncued rGT, stimulated females did not differ from controls in the number of infusions of cocaine taken (F_1,14_ = 2.39, p = 0.11), **(B)** the number of active responses (F_1,14_ = 2.39, p = 0.11), or the number inactive responses (F_1,14_ = 5.76, p = 0.82). **(C)** Stimulated males trained in the uncued rGT did not differ from controls in the number of infusions of cocaine taken (F_1,14_ = 0.19, p = 0.67), **(D)** the number of responses made on the active lever (F_1,14_ = 0.29, p = 0.60), or the number of responses made on the inactive lever (F_1,14_ = 0.29, p = 0.60). **(E)** Stimulated females trained in the cued rGT took more infusions of cocaine than controls (F_1,24_ = 15.87, p < 0.01), **(F)** made more responses on the active lever (F_1,24_ = 13.09, p < 0.01) but did not differ in inactive responses (F_1,24_ = 3.49, p < 0.07). **(G)** In the cued rGT, stimulated males took more infusions of cocaine (F_1,20_ = 7.60, p = 0.01) and **(H)** made more responses on the active lever (F_1,20_ = 9.64, p = 0.01) but did not differ in inactive responding (F_1,20_ = 0.63, p = 0.44). **(I)** Regardless of sex, C control animals trained in the cued rGT took more infusions of cocaine than those trained in the uncued rGT (F_1,36_ = 4.52, p = 0.041)

#### Premature responses

Cocaine self-administration increased premature responding in stimulated females trained in the cued rGT (session x task: F_14,994_ = 2.82, p < 0.01; session x group: F_14,994_ = 3.34, p < 0.01; sex: F_1,71_ = 4.83, p = 0.03; t_13_ = -1.79, p = 0.05) and effected a trend increase in stimulated females in the uncued rGT (t_7_ = -1.76, p = 0.06) but impacted the variable in no other condition (all t < 1.42, all p > 0.09) (Supplemental Figure 3A-D).

Cocaine self-administration decreased the number of trials performed by control females trained in the uncued rGT (session x group x sex x task: F_14,994_ = 1.82, p = 0.03; t_13_ = -1.79, p = 0.05) but impacted the variable in no other condition (all t < 1.42, all p > 0.09) (Supplemental Figure 3E-H).

Cocaine self-administration increased omissions in stimulated males in the cued rGT (session x sex s task x group: F_14, 994_ = 2.39, p < 0.01; t_10_ = -1.184, p = 0.04) but affected the variable in no other conditions (all t > 1.21, all p > 0.13) (Supplemental Figure 3I-L).

Cocaine self-administration had no effect on choice latencies (all F < 1.32, all p > 0.16; Supplemental Figure 3M-P) or collection latencies (all F < 1.17, all p > 0.29; Supplemental Figure 3Q-T).

**Supplemental Figure 3.**
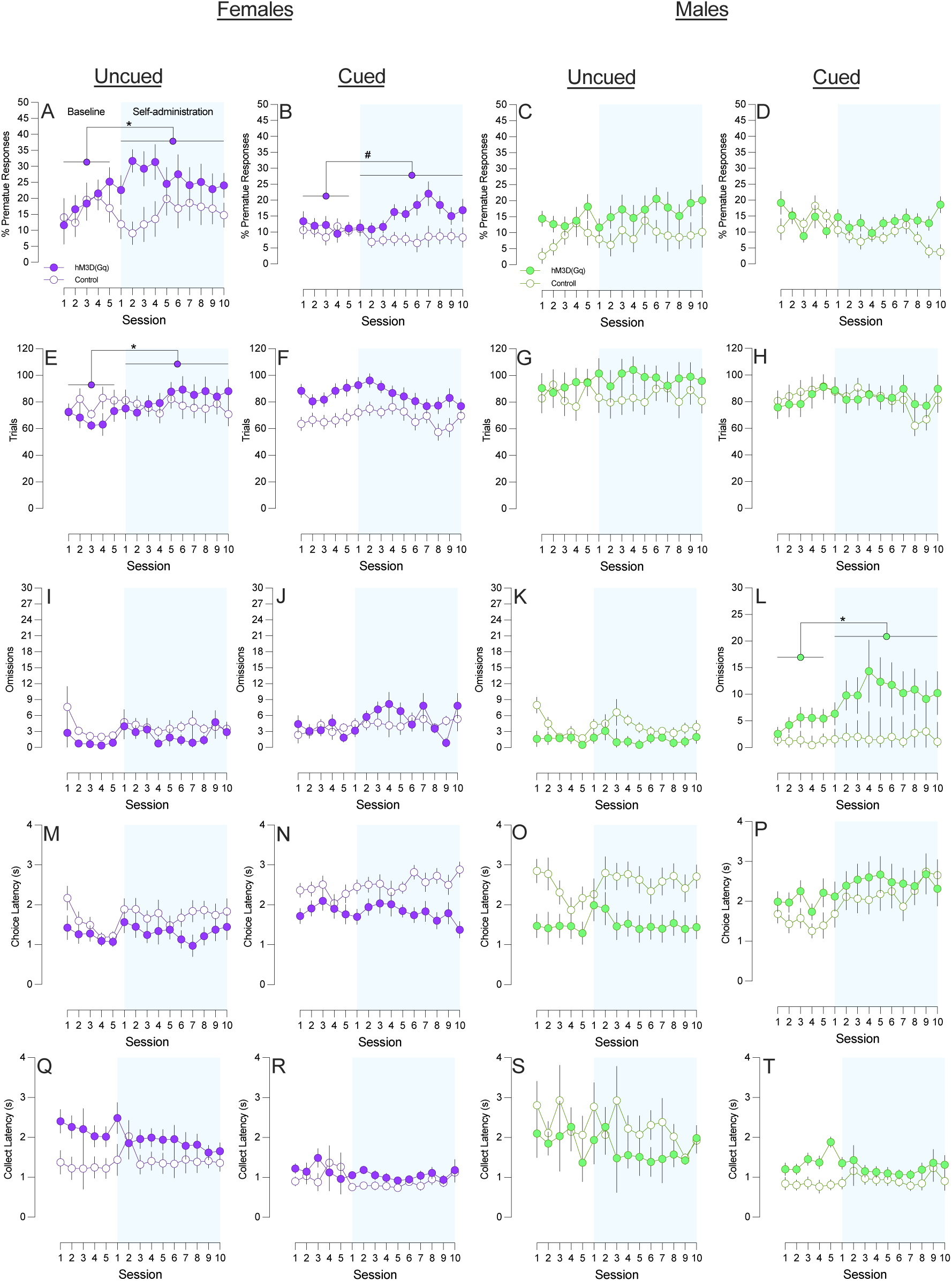
Effect of VTA DA neuron stimulation on cocaine-induced changes in auxiliary behavioural measures. **(A)** In the uncued rGT and **(B)** cued rGT, stimulated females exhibited increased premature responding following cocaine self-administration. **(C-D)** Cocaine self-administration did not impact premature responding in males in either task. **(E)** In the uncued rGT, stimulated females completed more trials following cocaine self-administration but **(F)** showed no differences in the uncued rGT. **(G-H)** Cocaine self-administration did not affect the number of trials completed by males in either task variant. **(I-J)** Cocaine self-administration did not impact omissions in females trained in either task variant. **(K)** Cocaine self-administration did not affect omissions in males trained in the uncued rGT but **(L)** increased omissions in stimulated males trained in the cued rGT. **(M-P)** Cocaine self-administration did not affect choice latencies or **(Q-T)** collection latencies. Error bars represent the SEM. “*” indicates p < 0.05. “**” indicates p < 0.01.

## Acknowledgements

This work was supported by a Canadian Institutes of Health Research project grant awarded to CAW (PJT-162312). TJH & MLM were supported by a Marshall Graduate Scholarship. CSC was supported by a Canadian Graduate Scholarship (CGS-M). This work took place at a UBC campus situated on the traditional, ancestral, and unceded land of the xʷməθkʷəy̓əm (Musqueam), sə̓lílwətaʔɬSelilwitulh (Tsleil-Waututh) and Sḵwx̱wú7mesh (Squamish) Peoples. We acknowledge and are grateful for their stewardship of this land for thousands of years.

## Notes

### Competing Interest Statement

The authors have declared no competing interest.

